# HistoMIL: a Python package for training Multiple Instance Learning models on histopathology slides

**DOI:** 10.1101/2023.06.02.543494

**Authors:** Shi Pan, Maria Secrier

## Abstract

Haematoxilin and eosin (H&E) stained slides are commonly used as the gold standard for disease diagnosis. Remarkable progress in the deep learning field in recent years has enabled the detection of complex molecular patterns within such histopathology slides, suggesting automated approaches could help inform pathologists’ decisions. In this context, Multiple Instance Learning (MIL) algorithms have been shown to outperform Transfer Learning (TL) based methods for a variety of tasks. However, there is still a considerable complexity to implementing and using such methods for computational biology research and clinical practice. We introduce HistoMIL, a Python package designed to simplify the implementation, training, and inference process of MIL-based algorithms for computational pathologists and biomedical researchers. In HistoMIL, we have integrated a self-supervised learning-based module to train the feature encoder, a full pipeline encompassing TL as well as three MIL algorithms, namely ABMIL (1), DSMIL (2), and TransMIL (3). By utilising the PyTorch Lightning framework (4), HistoMIL enables effortless customization of training intricacies and implementation of novel algorithms. We illustrate the capabilities of HistoMIL by building predictive models for 2,487 cancer hallmark genes on breast cancer histology slides from The Cancer Genome Atlas, on which we demonstrate AUROC performances of up to 85%. Cell proliferation processes were most easily detected, shedding light on the opportunities but also limitations of applying deep learning for gene expression detection. The HistoMIL package is proposed as a tool to simplify the implementation and usage of deep learning tasks for researchers.

## INTRODUCTION

Histopathology slides stained with haematoxylin and eosin (H&E) are widely considered as the gold standard for the diagnosis of cancer and other diseases. Deep learning (DL) (5) based approaches have demonstrated tremendous potential for reproducing the workflows of human experts employing such slides in a variety of tasks, e.g. diagnosing cancer and classifying tumour types (6, 7); segmenting sub-regions at the pixel level to identify nuclei or tissue boundaries (8, 9); and predicting important clinical metrics such as survival (10), recurrence rates (11, 12) and response to treatment (13). Several studies have shown that DL-based approaches can also help to predict more complex molecular labels that from Whole Slide Images (WSI) datasets, such as microsatellite instability, DNA damage repair deficiencies, mutations or expression of different genes (14-17). In particular, some studies have introduced MIL algorithms which outperform Transfer Learning (TL) based pipelines in different tasks such as survival prediction (18, 19). However, there are considerable complexities of implementing an MIL-based pipeline to predict molecular labels from WSI datasets.

Digital pathology WSI datasets include multiple large images scanned from original diagnostic slides stained with H&E. Each of these images normally contain giga-pixels in a single file. This brings about specific challenges when applying MIL-based methods: (1) WSI files cannot be directly read by widely used image processing packages such as PIL (20), (2) classic architectures of a neural network are designed for lower resolution (i.e. 224×224 pixels (21)), and (3) loading entire a batch of WSIs during training is almost unmanageable and untraceable due to the limited GPU memory. There are various approaches to tackle these issues, and from the perspective of toolkit design, they can be roughly divided into: slide reading packages, pre-processing packages, and machine learning (ML) oriented packages. The earlier implementations primarily focused on designing a user-friendly API to read WSIs, while an increasing number of packages are now being designed to meet the demands of machine learning algorithms.

### WSI reading packages/libraries

such as OpenSlide (22), BioFormats (23), HighDicom (24), are typical tools that provide a user-friendly API to handle WSI data formats. Researchers can access the raw data of various WSI files by using these libraries. However, these packages are not designed for training neural networks. OpenSlide includes a Python interface but does not support some formats (i.e. OME-TIFF format) (22). BioFormats (23) and QuPath (24) support more WSI formats. QuPath also has a graphical user interface which makes it easy to use for beginners. However, both of them rely heavily on some Java libraries which make them potentially difficult to integrate with Python-based workflows and can be very slow in some extreme cases. As a bioimage analysis program with well-supported documentation and Python interfaces, QuPath allows users to access packages (such as scikit-learn) to implement a simple ML pipeline (24), which has been a milestone in the development of WSI processing toolkits. However, this tool is not designed for DL algorithms or MIL algorithms.

### Pre-processing packages

have also attracted much attention in the past years. While some algorithms for WSI classification tasks include similar basic components, e.g. WSI reading, patch extraction and colour normalisation (25, 26), a comprehensive and broadly applicable ML-oriented package is still very difficult to design. The patching step, which consists of dividing a whole slide into smaller tiles (or “patches”) that can be more effectively processed individually, is a good example to demonstrate the difficulties of using existing pre-processing packages. Patching is a memory-expensive and time-consuming step in the preprocessing stage. While scale-pyramid and multi-thread processing strategies can be effective in this instance (27), most pre-processing packages do not implement them. In packages such as CLAM (28), developers have incorporated a comprehensive preprocessing workflow with high-quality code. But these packages prioritise the implementation of a particular algorithm, which could present challenges when attempting to utilise their preprocessing steps with other algorithms. Semi-automated packages such as PyHIST (29), Deep-histopath (30) and ASAP (31) normally include a focus on the tissue segmentation or patching step. However, some of them are only designed for particular datasets, e.g. Deep-Histopath is designed for The Tumour Proliferation Assessment Challenge 2016 (TUPAC16) (30).

### Machine Learning oriented packages

are designed for training models on WSI datasets. Some open source packages such as DeepMed (26) or TIAToolBox (27) aim to provide a common platform of training deep learning models, and simplify the inference step of pre-trained models. We emphasise that these packages are different from pre-processing packages due to the support provided for training machine learning or deep learning models. Nevertheless, these packages have their own limitations. Stain normalisation, a technique employed to ensure slides with different colour ranges are transformed in a uniform way to enable comparisons, can be a good example to demonstrate the limitation of the packages that only consider one preprocessing step. Packages like stainlib (25) work well for the colour normalisation step, but require extra effort to transfer WSI raw data into a readable format such as JPEG or PNG. ML-oriented packages can handle the entire pipeline which generally starts by reading WSI raw data and ends with the model training/inferencing step. Moreover, packages such as PathML (32) aim to deliver richer content for their users by integrating more extensive models such as a tissue segmentation model. Also, researchers can easily and quickly train DL models on WSI datasets by using the TL protocol. For some of the open source packages, various useful tools have been integrated together, e.g. cell segmentation (32) or graph aggregator (27). Packages such as TIAToolBox also offer improved scalability through their module design and unit-testable code (27). However, the existing pipelines are not fully utilising the capabilities of deep learning approaches.

One of the major challenges is that packages such as DeepMed only support TL-based models (26). Oftentimes, researchers can only access slide-level or patient-level annotations, as obtaining pixel-level annotations can be prohibitively expensive or impractical. Existing transfer learning-based approaches only consider the original slide-level or patient-level labels, assigning them to each partitioned patch as training label information. In thise TL-based protocol, introducing pseudo-labels can be considered a necessary step when pixel level labels are not available. This, however, adds more noise to the model training process. Training a batch of models may also be a problem for packages like PathML (32). The target model may learn misleading information from pseudo-labels and be stuck in a local optimal point. In extreme cases, for instance, the areas containing tumours may be exceedingly minute, with merely a handful of patches embodying genuine tumour information. Under such circumstances, assigning identical labels to all patches would evidently interfere with model learning, and even algorithms based on attention mechanisms may be impacted by such extreme cases, thereby affecting the model’s generalisation capabilities (1). As another example, a tumour area could be labelled as highly expressing a specific gene that is only expressed in a subtype of T cells, and in this case the target model may learn complex distributions for all cells populations within the tumour tissue and incorrectly classify all cells as positive cases.

In practice, researchers may want to have a general understanding about how multiple biomarkers are spatially distributed by predicting those molecular labels from H&E slides. They may want to predict hundreds of target labels in an initial setting. Existing packages cannot enable researchers to quickly scale-up an algorithm for multiple molecular labels, as most existing packages are not designed to deploy a batch of models. Additionally, existing libraries frequently use pre-trained models based on ImageNet as feature extractors. Although some recent work demonstrates that feature extractor networks which are trained by self-supervised learning protocols on WSIs may increase the final performance (2, 18), training feature extractor networks with SSL protocols is not supported by existing ML-oriented packages.

To address some of these challenges, we introduce HistoMIL, a new DL package based on PyTorch (33) and PyTorch-lightning (4) that can simplify the training of WSI-based classification models. This package provides a complete pre-processing pipeline that simplifies the pre-processing stage and decreases the complexity of converting raw WSI data into usable data for DL frameworks. We implement multiple MIL models that allow flexible training against different target labels, along with multiple SSL models that simplify the training of feature encoders. We demonstrate the performance of HistoMIL on predicting >2,000 cancer-relevant molecular labels within breast cancer H&E slides from the Cancer Genome Atlas (TCGA), and highlight specific pathways whose activity can be reasonably captured within histopathology tissue.

## METHODS

### Overview of the HistoMIL package

The HistoMIL library includes three levels: **data, model**, and **experiment** (**Figure 1a**). The **data** level includes several data pre-processing steps (WSI reading, tissue segmentation and patching). Similarly to other ML-oriented packages, we include image normalization functions and feature extraction to save intermediate data and accelerate model training. Additionally, we introduce a cohort level to handle metadata such as patient information, molecular labels, and other additional information. The design of HistoMIL follows the evolution of relevant packages in literature (**Figure 1b**).

**Figure 1:**
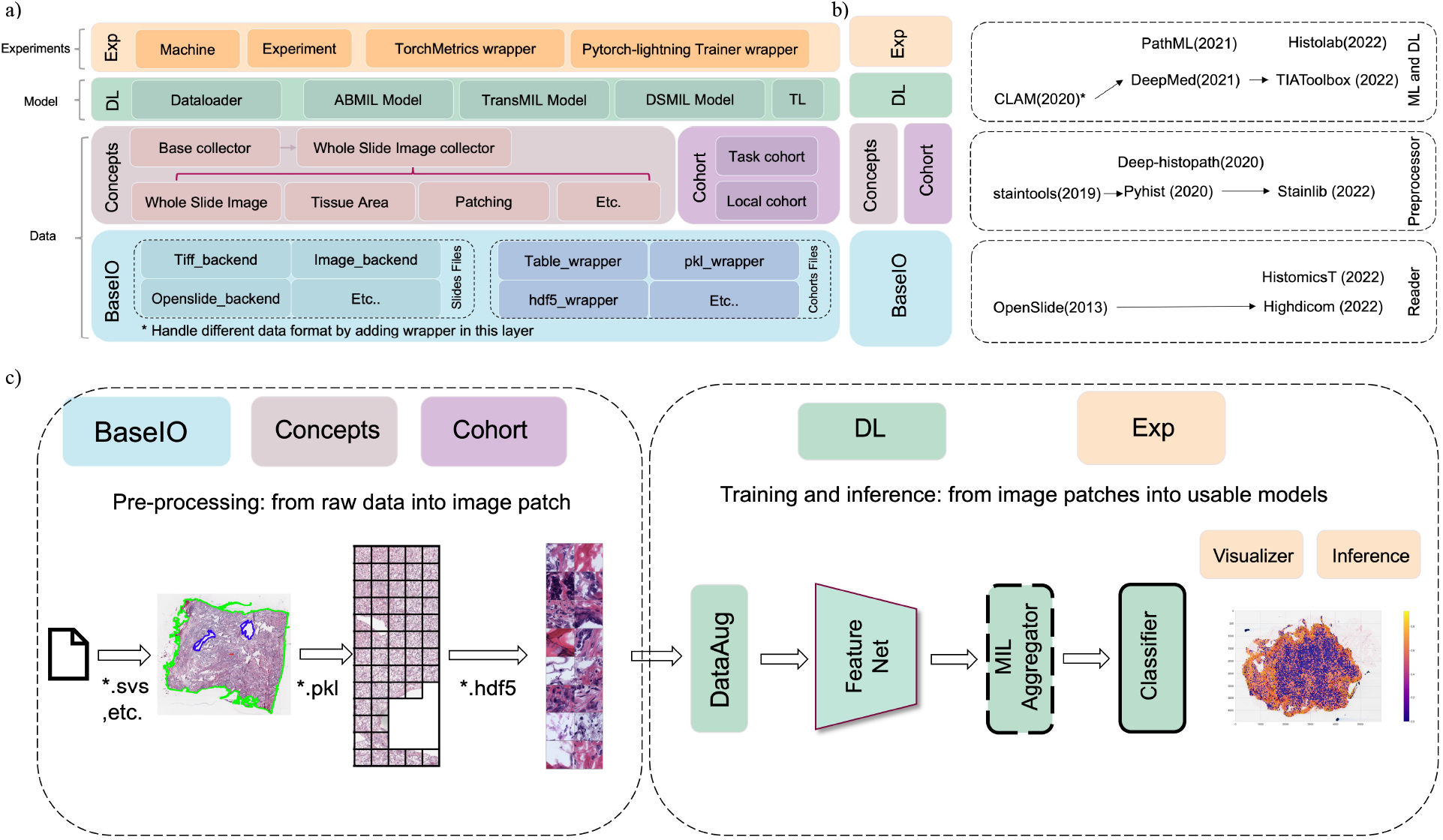
Overview of HistoMIL package design and working pipeline. (a) The diagram illustrates the relevant modules comprising HistoMIL and the logical structure organizing them. HistoMIL features four levels encompassing file reading, WSI-related processing modules, deep learning algorithms, and experiment management modules. (b) The relationship between HistoMIL modules and state-of-the-art libraries. The proposed package covers the entire workflow of WSI reading, preprocessing, and MIL algorithms, a design informed by the primary functionalities of other packages in the literature. (c) How HistoMIL’s various modules operate within an actual pipeline. We enumerate a typical processing workflow, including preprocessing, MIL training, and inference components. It is evident that different modules correspond to distinct functional parts within the process.

The **model** level can be split into two sections: the backbone part and the MIL method. By integrating the interface of the timm PyTorch image model (33), our backbone module can download various backbone network architectures and pre-trained parameters for feature extraction. Additionally, our package allows users to apply self-supervised learning protocols to train the feature extractor from scratch by only using WSI datasets. Multiple MIL methods and a baseline TL model are included in our package. The entire algorithm implementation is based on PyTorch-Lightning (4), which makes it possible to quickly train and fine-tune models for a new label. Furthermore, HistoMIL includes pre-defined parameters as default settings, which allows users to quickly try out and scale-up an algorithm for different targets.

At the **experiment** level, setting up different models for various datasets and hardware conditions can be done easily. Researchers can simply initialise a series of instances to search over the hyperparameter space defined by the related customisable model *para* class. An instance of *Trainer* class will initialise a PyTorch lightning module to automatically fit the requests of hardware. Meanwhile third-party tuners such as the Ray tuner (34) can directly use those module instances, offering us the capability to find the optimal model configuration easily. **Figure 1c** demonstrates which modules are used in different steps of a process that employs a MIL algorithm. The details about how modules interact with each other during preprocessing, self-supervised learning and MIL can be found in **Figure 2**.

**Figure 2:**
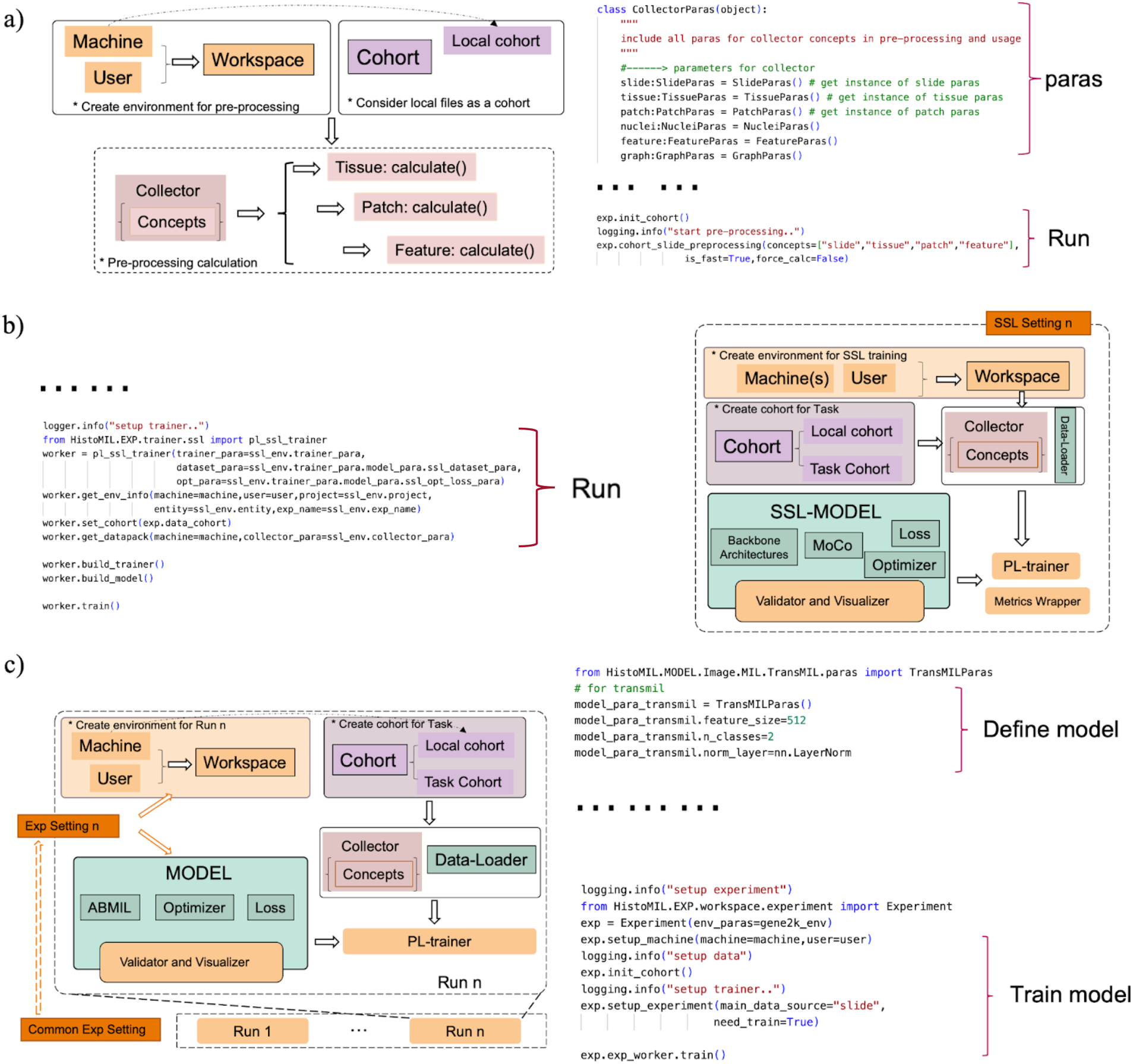
Diagram depicting three primary application scenarios for HistoMIL and their corresponding code blocks. (a) The invocation method for HistoMIL during batch preprocessing. With predefined processing parameters, HistoMIL can process an entire dataset in one go using just three to four simple commands. (b) The code and calling logic for SSL training. HistoMIL facilitates SSL on WSI datasets by predefining SSL-related parameters and modifying the trainer accordingly. (c) The code and module calling process for MIL training. Compared to SSL, MIL adds adjustments to the model’s parameters while simplifying the training setup process. For users with access to a GPU server, numerous models can be trained simultaneously by configuring parameters in the bash file.

### Data level design

The HistoMIL design includes a data level to handle different data formats, pre-processing and other meta-information of the target cohort. Pre-processing WSIs can be time-consuming and computationally expensive in many cases. By integrating a number of configurations and functions, the HistoMIL package offers a functional interface to smoothly run each necessary step in a different pipeline, and save the intermediate output as needed. We divide the pre-processing steps into three separate concept categories: tissue, patch, and feature. Each of these concepts is linked with a parameter class to help users modify the pre-processing steps by following their own experimental requirements. A manager class named *WSI_collector* helps users unify the initialisation, calculation and loading process of these five concepts to further decrease the complexity of usage. Also, the package offers several designs to further accelerate these steps.

#### Reading Raw WSI data

HistoMIL can handle different data formats by implementing slide_backend wrappers for widely considered packages such as OpenSlide-Python (22). This design can help researchers access the raw data of different WSI formats and provides a common interface for the subsequent steps. For instance, the scale pyramid in WSI processing can be more important than other areas as some WSI files naturally include multi-scale representation of the raw data (27). A series of downscaled or upscaled versions of the original slide will help the potential deep learning models get more information. HistoMIL will create a common scale pyramid as metadata of a WSI file. Also, researchers can implement customised slide wrappers to access additional slide formats. This modular design enables the rapid integration of customised slide backends into existing pipelines, thereby facilitating the support of different datasets.

#### Tissue masking and patch extraction

WSIs typically involve a considerable area that only includes non-biologically relevant background elements such as glass or marker pen. HistoMIL includes a *Tissue* class to identify and remove these areas by generating a tissue mask. Inspired by other ML-oriented packages, the *Tissue* class includes a wrapper for an Otsu function to categorise pixels into foreground or background (28). Some basic morphological operations are integrated to eliminate small holes within the tissue region. Researchers can also implement different wrapper functions for different purposes. Patch extraction, which aims to decrease the cost of GPU memory when training a model, works in a similar manner. A large WSI will be partitioned into small patches by using an integrated function to iteratively walk through the tissue area. All the intermediate data will be stored for further usage, and the proposed implementation can avoid filling available memory with enhanced memory efficiency. By using this scale pyramid, we configure the segmentation process in higher scale pyramid levels as our default setting to decrease computational costs. Similarly, the default settings of patch selection functions, which can decide whether current patches need to be processed in a pipeline, are applied at a high pyramid level. We use a simple multiple processing pool to further accelerate the patching step, similarly to the CLAM package (28). All these parameters (such as step size and window size in the patch concept class) can be easily modified in related parameter classes to fit different requests.

#### Feature Extraction

A pre-calculation step for the feature extraction part may decrease time costs during training. Some MIL algorithms (2, 18) (3) can work with pre-trained feature extraction networks and choose to calculate feature vectors for each selected patch in a pre-processing step. By following this paradigm, we designed a *Feature* class to synchronise the manipulation and maintenance of the feature vectors generated from each slide. This design can also simplify the potential clustering process for feature vectors within each bag or clustering on the entire dataset. Hence, preprocessing, which would have required the implementation of multiple functions and methods, has been simplified to a few lines of code invocation in HistoMIL (Figure **2a**).

### Model training

In HistoMIL, model training is a crucial part especially for the target molecular labels that may be difficult to identify by only considering tissue morphology differences. MIL has been introduced as a higher performing weakly supervised learning protocol for WSI classification. A slide (or WSI) is considered as a bag of instances in the MIL protocol. Each bag may contain hundreds or thousands of instances (patches) that are assembled into slide-level features for classification. To the best of our knowledge, no existing package to date has been designed to handle MIL methods. There is considerable complexity in building a MIL pipeline for WSI classification tasks due to the inherent complexity of WSI formats, heterogeneous nature of cell morphology, and unbalanced target labels. HistoMIL is designed to simplify the implementation of MIL-based pipelines by implementing Self-Supervised Learning and Multiple Instance Learning modules (**Figures 2b-c**).

#### Self-supervised learning for feature extraction

End-to-End training for a MIL pipeline that consists of feature extractors and aggregators may be computationally expensive, especially in a large WSI. Therefore, the existing model either uses fixed patch features derived from CNNs or simply employs a small number of high-scoring patches to update feature extractors. Inspired by the TL paradigm, the feature extractor network can re-use pre-trained parameters from general image classification domains (e.g. ImageNet). The advantage is that there are different pre-trained feature extraction networks that can be chosen for different tasks. Thus, target classification models can be trained quickly with a pre-trained feature extractor network. However, if a pre-trained feature network is not trained on a WSI dataset, it may require extra effort in the pre-processing steps and the model may not converge. The process of aggregating and classifying may also be susceptible to the issue of overfitting and inadequate supervision, resulting in potential bias and other limitations.

Self-Supervised Learning (SSL) is frequently mentioned as a solution for the mentioned problems. While some studies, such as (28), have shown that the SSL phase is a crucial element in a pipeline, integrating the WSI dataset into an SSL pipeline remains difficult. HistoMIL offers an easy way to accomplish this part by using a user-friendly SSL module. To train a feature extractor, the HistoMIL package offers an easy way to apply self-supervised training methods. Firstly, we introduce a trainer class to simplify the training process on a WSI dataset. A wrapper class is created for the timm package (33) which includes most of the popular backbone architectures and potential pre-trained parameters. Some widely employed SSL methods have been implemented, such as the MoCo (35, 36) and SimCLR (37) methods. By using these built-in methods, HistoMIL also offers predetermined hyperparameters to further simplify the training process to new users. Also, it is specifically designed for WSI data. Users can simply download raw data from projects such as TCGA and do not need to worry about pre-processing and data augmentation. In addition, utilising the feature extractor network trained by HistoMIL’s SSL module negates the necessity to account for discrepancies in operator implementation that may arise when importing trained parameters from other packages. Moreover, by embedding SSL methods within the HistoMIL package, researchers can effortlessly incorporate the selection of SSL methods and backbone architectures as hyperparameters when optimising their models.

#### MIL methods

To train a slide-level classifier on target labels, HistoMIL offers a variety of algorithms that are ready to use. To the best of our knowledge, no existing package can simplify the training and classification process for MIL algorithms in WSI datasets. Transfer Learning (TL) protocols and Multiple Instance Learning (MIL) protocols have both been widely considered by researchers for classification tasks on WSIs. While packages such as DeepMed (26) have been designed to apply TL-based models on WSI datasets, there are considerable difficulties in implementing a MIL model for WSI classification tasks. Whole slide images often encompass several gigabytes or even terabytes, and the aggregation function of MIL methods may suffer due to its high-dimensional feature space, with tens of thousands of feature vectors per slide. Moreover, there is the lack of a standard implementation for various MIL algorithms. For molecular labels, training a MIL model may be even more difficult when faced with problems such as heterogeneity of cell morphology and imbalanced data. In some extreme cases, the target classification model may be vulnerable to overfitting, and is unable to explore rich feature representations because of the insufficient supervised signal. All of these challenges require users to have considerable experience with implementing and training deep neural networks. This requirement may exclude researchers who need these models but lack the necessary experience. HistoMIL simplifies this aspect by introducing built-in MIL modules with pre-defined hyperparameters.

In HistoMIL, a customizable sampler function in our Dataloader implementation is introduced that only samples some instances from each bag. Different batch sizes can be used if the MIL algorithm needs to sample N instances from each bag. But if all the patches must be read in at once, the batch size should be fixed at 1 and this may lead to an unstable training process. In our default setting, we chose to decrease the initial learning rate and accumulate gradients over the training steps to smooth the optimisation process. We also include a cohort instance during training to handle patient metadata, which means users can also assign pseudo labels for each patch based on the label per slide, and this enables users to train their model using the transfer learning paradigm as well.

### Trainer and experiment setup

A significant amount of time and effort is spent on the tuning process and hyperparameter selection steps when training on WSI datasets. We designed the *trainer* class and *experiment* class in HistoMIL, which are built on the PyTorch Lightning framework, to cut these costs. Our target is to simplify the tuning process for different target labels. All of the SSL, TL and MIL algorithms mentioned above are implemented using a related PyTorch Lightning module in HistoMIL. Researchers can initialise a PyTorch Lightning instance with HistoMIL, and this instance can be easily fed into third-party tuners such as the Ray tuner package (34).

### Experimental setting and dataset

In this paper, we exemplify the power and versatility of HistoMIL in oncology-related tasks by building prediction models for 2,487 cancer-related genes.

#### Dataset

The Cancer Genome Atlas (TCGA) is a collaborative project that aims to characterise the genomic and molecular landscape of various cancers (38). We chose TCGA-BRCA as the largest available dataset with diagnostic H&E-stained slides and matched RNA-sequencing which we could use to define cancer-relevant labels (n=2,487). The TCGA-BRCA dataset contains a large amount of data, including whole slide images, genomic data, clinical data, and more. It includes samples from a diverse patient population, including patients of different ages, races, and ethnicities. This helps avoid biases as potential factors that may affect model performance. As a widely used data source, the TCGA-BRCA dataset has undergone quality control, which ensures that the data is of high quality and well-annotated. This can help reduce the noise and variability in the dataset. We downloaded 1,133 diagnostic WSIs from the Genomic Data Commons (GDC) Data Portal (39). In our experiment, we split WSIs into patches, then automatically selected WSIs with more than 1,000 patches, and further removed the images which were blurry or containing marks. This left us with 1,012 WSIs for analyses, which we split into 80% for training and 20% for testing.

#### Prediction task

The target labels for the experiments were extracted from the RNA-seq profiles of the same TCGA-BRCA patients for which H&E-stained diagnostic slides were also available, downloaded using the TCGAbiolinks package (40). We surveyed 2,487 genes involved in various cancer hallmark pathways derived from MSigDB (41) using the msigdbr R package (see **Supplementary Table 1**), and used the FPKM normalised expression values for each gene to categorise tumours as “highly” or “lowly” expressing the respective gene based on the median split. The HistoMIL package was used to train and validate prediction models for the expression of the selected 2,487 cancer hallmark genes. Functional enrichment analysis was performed using GeneMania (42). By predicting the expression of a gene of interest throughout the entire tissue slide, researchers can highlight the features derived from the attention score or gradient vectors. This could aid in diagnosis or help pathologists identify subtle differences in gene activity that might not be apparent in traditional histological sections. Extracting features and analysing results is normally a time-consuming and computationally intensive task. By using HistoMIL, we can train models on a very large scale. This may lead to faster and more efficient analysis of whole slide images in future clinical usage.

### Software and package

HistoMIL utilises Python as the primary implementation language and PyTorch as the underlying deep learning platform (33). Using these libraries, HistoMIL can easily implement customised algorithms. Furthermore, we introduced PyTorch Lightning as a higher-level framework to simplify the implementation of training code (4). All the built-in algorithms in HistoMIL adhere to PyTorch Lightning’s design philosophy, which decomposes the training process into different functions. This facilitates user customization while also ensuring concise implementation. The HistoMIL package is available at the following GitHub repository: https://github.com/secrierlab/HistoMIL.

## RESULTS

In this section, we demonstrate the usability and scalability of HistoMIL in WSI-level prediction tasks using recently proposed deep learning DL models. One clinically relevant task in cancer that has been recently made possible by modern computational pathology is assessing the state of specific genes or even entire molecular pathways directly within the histopathology tissue slide using deep learning techniques (43, 44). Such rapid assessment can assist diagnosis or inform prognosis and treatment. Typically, this evaluation is completed through gene panel profiling or immunohistochemistry (IHC). However, these tests can result in time delays and additional costs as they are used as additional steps after visual inspection of routinely stained H&E sections. HistoMIL simplifies this task by enabling fast and efficient implementation for predicting diverse molecular labels in H&E-stained slides.

Here, we demonstrate the potential of HistoMIL to accomplish all the necessary steps for predicting thousands of cancer hallmarks, while simplifying the entire analysis workflow. We used 1,0133 H&E-stained slides of breast cancer tissue and matched RNA-seq data from TCGA to train over 8,000 models for the classification of 2,487 cancer hallmark genes. First, we processed the WSI dataset using pre-defined pre-processing functions. This step involves extracting the tissue area from H&E histological images, generating patches automatically, and saving them as H5DF and image files. Subsequently, we trained the Feature Extractor Network using the SSL module and predicted the target gene expression labels using the built-in MIL algorithms. As depicted in **Figure 3a**, the implementation of the pipeline is simplified through the use of the generic interface provided by HistoMIL. This reduces the effort required by new researchers seeking to expand these methods.

**Figure 3:**
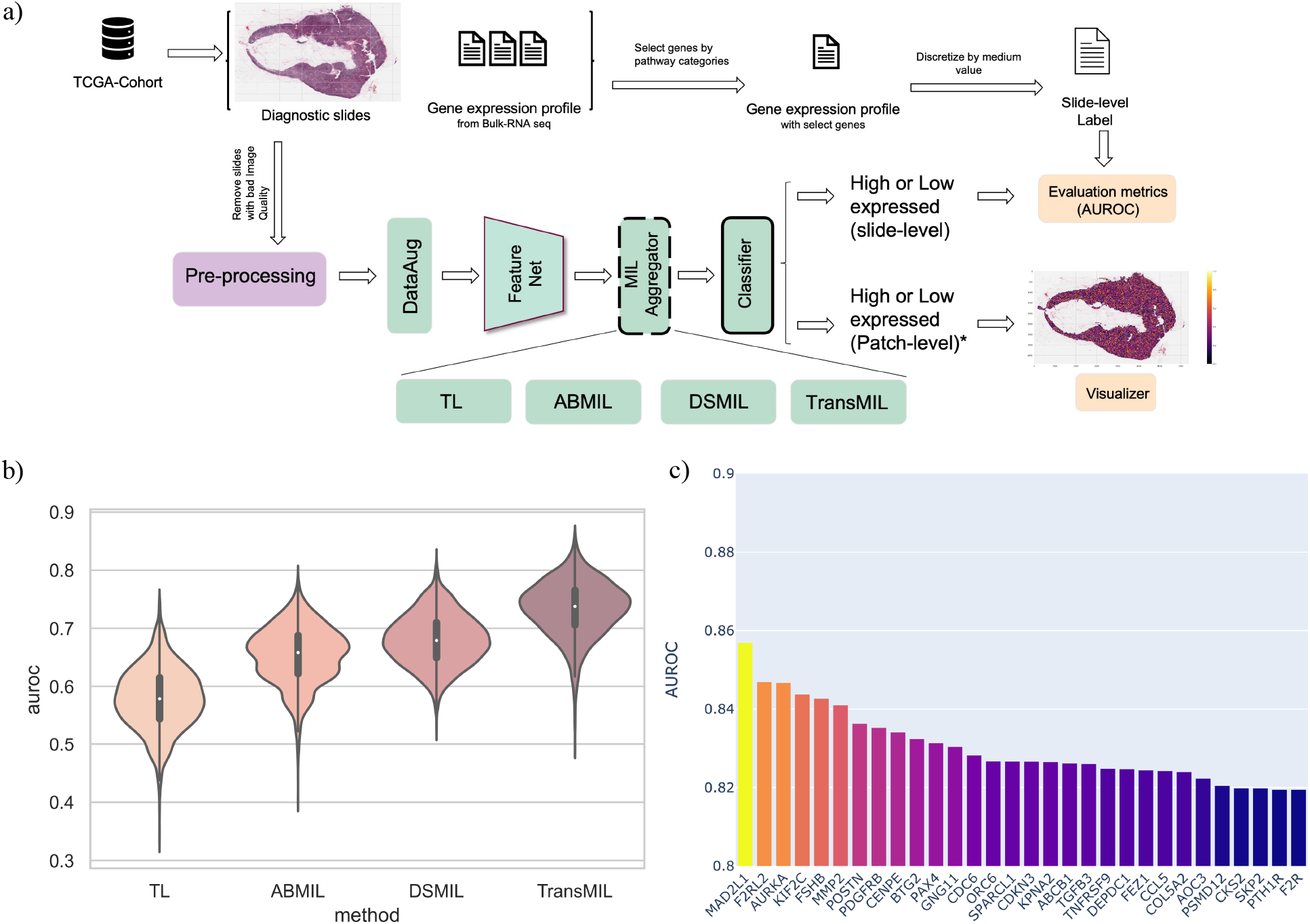
Experimental workflow and performance comparison. (a) The diagram showcases the complete workflow for utilizing HistoMIL in predictive experiments. TCGA-BRCA’s raw data undergoes preprocessing, yielding image patches and associated labels suitable for MIL processing. The experimental task involves predicting gene expression, with the model’s performance displayed as AUROC. (b) Comparison of performance distribution among different algorithms in predicting the expression of 2,487 cancer-related genes. Each distribution contains 2,487 data points. TransMIL exhibits superior performance relative to the other algorithms. (c) The top 30 genes with the highest AUROC scores. In the test set, the model’s accuracy in predicting gene expression levels (high or low) reaches up to ∼86%.

Despite the evident advantages of using MIL algorithms for H&E-based predictions, researchers without much coding experience may face challenges in replicating MIL pipelines, where slight variations in code may result in vastly different outcomes. Moreover, due to the complexity of handling high-dimensional histological data, novice researchers may be deterred from implementing this method. To train our models, we followed the same methodology as outlined in the original papers of TL (26), ABMIL (1), DSMIL(2), and TransMIL (3), and utilised the PyTorch Lightning wrapper designed by the HistoMIL library for the training steps. With the use of this package, these steps are easily accomplished, and the results are reproducible even without a a deep understanding of MIL algorithms. For each algorithm, we trained more than 2,000 models to predict expression levels for selected cancer hallmark genes.

Due to HistoMIL handling the complex WSI processing steps, the pipeline has been reproduced in an instance notebook with the toolbox as the backend. This can be found in the *Notebooks* folder of the HistoMIL GitHub repository (see Methods). We utilise the code snippets and fewer lines of code to demonstrate how to use HistoMIL. This highlights the effectiveness of the proposed package for WSI prediction tasks. Additionally, the implementation of the models includes options for calculating an attention score or providing patch level predictions, which allows MIL algorithms to predict the target labels for individual patches. In this example, we chose to focus on the WSI-level labels, with one expression value available per gene and per slide. To reduce the training and inference time, the example models in the package did not perform additional data augmentation, yet the models were still able to successfully predict the target label.

Predicting gene expression labels in the benchmark TCGA breast cancer dataset showcases the potential of using HistoMIL on WSI datasets. All of the predictions are based only on WSI data. The target labels are expression levels of 2,487 different cancer hallmark genes derived from MSigDB, selected as a benchmark task to demonstrate the scalability of our package. We considered this task a typical weakly supervised learning problem and selected the TransMIL method to learn the aggregation function. By using our EXP-level interface, we considered the ResNet-18 pre-trained feature extractor as a backbone network and all of the training processes were based on the PyTorch Lightning platform with early stopping.

Our experiments involved two training paradigms (TL and MIL) and a total of four different deep learning algorithms: TL, ABMIL (1), DSMIL (2), and TransMIL (3). From the experimental results, MIL algorithms generally outperformed TL algorithms when using the same feature encoder (**Figure 3b**). There were also performance differences among different MIL algorithms, but they generally followed the order of TransMIL > DSMIL > ABMIL. Considering the characteristics of these algorithms, it can be reasoned that the introduction of attention mechanisms and neighbourhood information have brought significant performance differences. Firstly, the slide-level attention mechanism could help the algorithm compare the importance among patches within a slide for classification. Additionally, TransMIL, which introduces neighbourhood information, has achieved the best results in our experiment. This is possibly due to the neighbour information capturing the broader changes in tissue structure which may be relevant in the context of a certain gene being expressed or deactivated.

Among the top genes where different models demonstrated good performance metrics on the test set (**Figure 3c, Supplementary Table 2**) were *MAD2L1, KIF2C* and *AURKA*. These are cell cycle regulators involved in spindle assembly and stabilization, and they promote chromosome segregation during mitosis (45-47). Other top-ranking genes such as *F2RL2* or *FSHB* are involved in G protein-coupled receptor signalling (47), while *MMP2* and *POSTN* are involved in matrix remodelling and cell adhesion (48, 49). These highly relevant cancer-promoting processes would be expected to leave a clear morphological trace in the tissue. Therefore, the activity of these genes might be more easily identifiable due to linked changes in tumour cell morphology and tissue structure as seen in the H&E slides. In addition, since different MIL algorithms show similar performance in predicting the expression of these genes, it further indicates that the expression patterns of these genes in H&E slides have a certain predictability. These patterns can be easily captured throughout the entire slide using heat map gradients of expression, thus informing on the spatial distribution of activity for a specific gene throughout the entire tissue (**Figure 4**).

**Figure 4:**
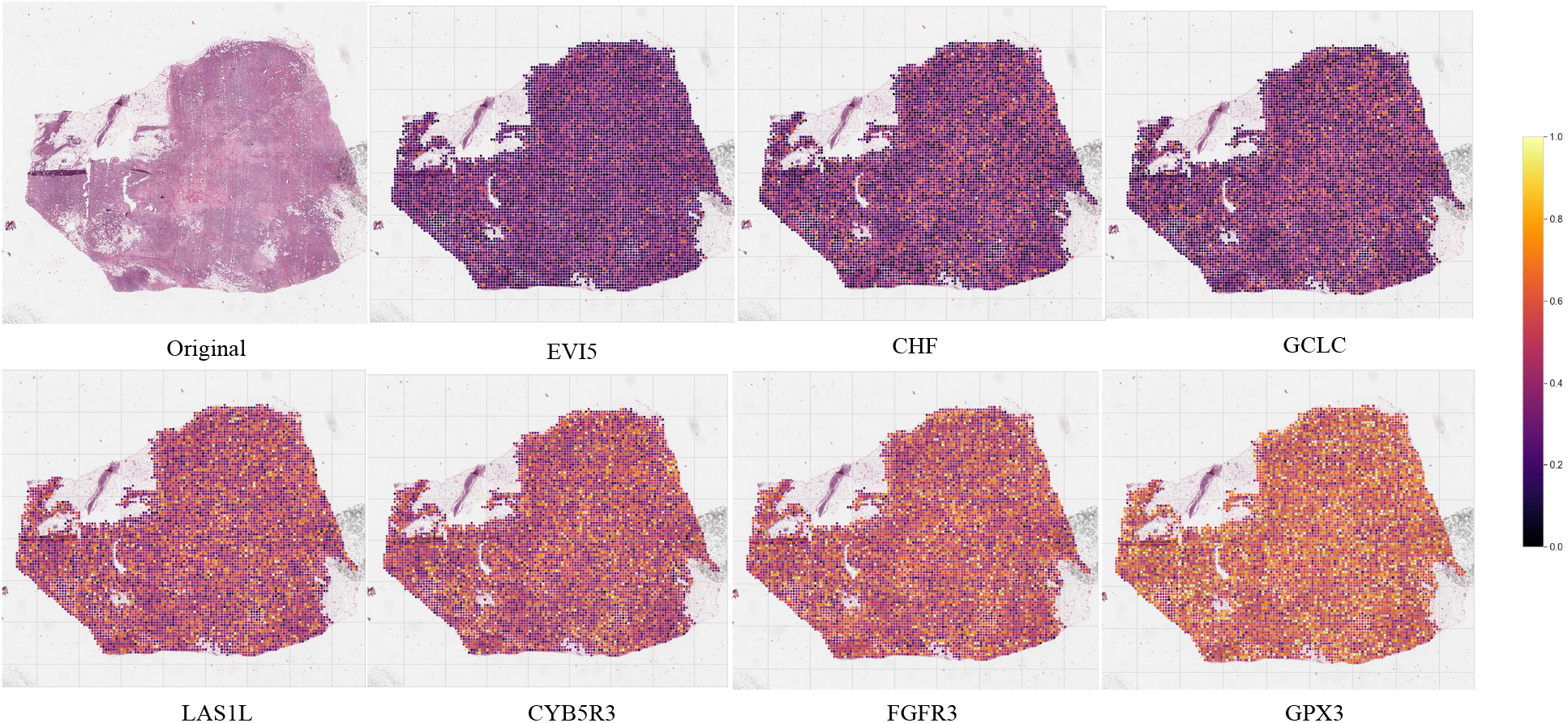
Model predictions for gene expression throughout the entire slide. The first image on the left in the top row is the original tissue slide. The remaining heat maps in the first row demonstrate the spatial distribution of gene activity for selected lowly expressed genes that were predicted by the model. The heat maps in the second row exhibit the spatial distribution of gene activity for highly expressed genes that were accurately predicted by the model. The purple to yellow gradient indicates increasing levels of predicted gene expression per patch. Intratumour heterogeneity of gene expression can be observed, particularly for *LAS1L* or *CYB5R3*.

Some genes, on the other hand, are more difficult to predict (**Supplementary Table 3**). For instance, the performance of the TransMIL algorithm is poor when predicting the expression levels of *SPRR3*, a marker for terminal squamous cell differentiation linked with tumour progression in early stage breast cancers (50), and *PGAM2*, a gene involved in oxidative stress responses (51). This could be explained by the complexity of the regulatory processes driven by these genes which may not render clear morphological changes in the cells. This could also be compounded by tumour heterogeneity, e.g. if oxidative stress is present in only part of the cancer tissue. Furthermore, unlike *MAD2L1, F2RL2*, and *KIF2C*, genes such as *SPRR3* and *PGAM2* exhibit higher performance variability among different MIL algorithms, and their performance is normally lower than an AUROC of 65%.

To further explore the capability of MIL methods, we focused on the higher-level activity captured by individual genes within the tissue, which can be summarized within hallmark pathways underlying cancer initiation and progression. These included processes such as *angiogenesis*, which plays a crucial role in the formation of new blood vessels to support tumour growth, *hypoxia*, triggered when cancer cells experience inadequate levels of oxygen, or the *P53 pathway*, which regulates cell cycle arrest and apoptosis in response to DNA damage (see **Supplementary Table 1** for a complete list). Across 14 key hallmark pathways we observed markedly consistent levels of performance of different MIL algorithms in predicting the expression of the genes involved in the respective pathways (**Figure 5**).

**Figure 5:**
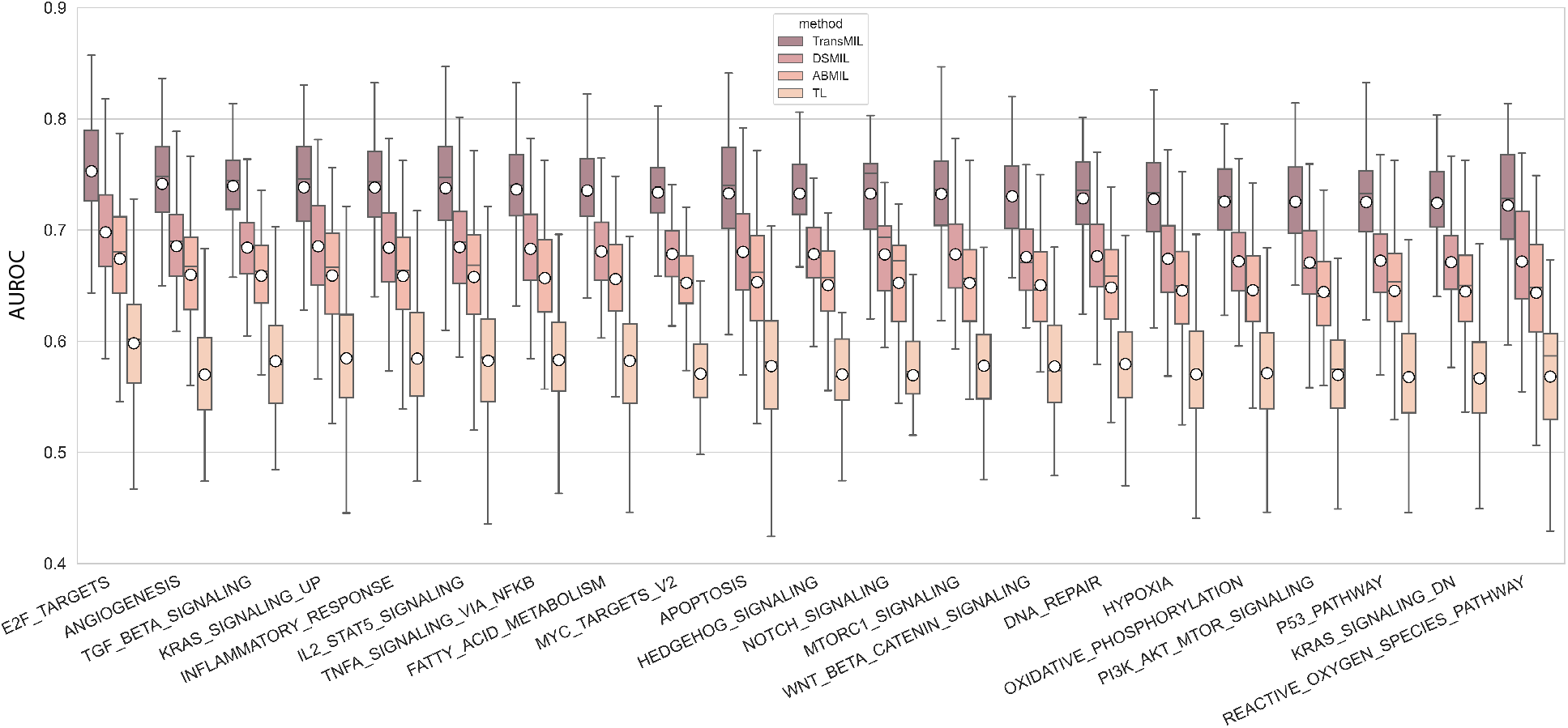
Performance of various algorithms on predicting pathway-level activity in cancer. TransMIL, DSMIL, ABMIL and TL (Transfer Learning) algorithm performance is compared across selected pathways. The boxes depict the AUROC distributions for each algorithm, coloured according to the legend. White circles represent the median values, while black lines indicate the mean. The different hallmark pathway groups are arranged from left to right in descending order of median values according to the TransMIL algorithm.

The E2F target genes had the highest average AUROC in our experiments (**Figure 5**). This pathway participates in the cell cycle G1/S transition and DNA replication, and is generally upregulated in tumour cells, leading to abnormal cell proliferation (52). Such proliferation differences may be more easily captured in the morphology of the cells as well as nuclear staining within H&E-stained sections. In fact, the top 50 highest performing models were for genes involved in cell cycle checkpoints, mitosis and DNA integrity (**Supplementary Table 4**), suggesting that cell division-related processes are most easily captured within cancer H&E slides using this methodology. This is not surprising, given the remarkably high performances (>90%) of deep learning models when it comes to distinguishing tumour areas from normal cells (53) considering that proliferation is the key defining hallmark of cancer. In contrast, the bottom 50 least performing models (with AUROCs < 62%) were for genes involved in tyrosine kinase and apoptotic signalling, nucleotide excision repair and oxidative stress responses (**Supplementary Table 3**), which might not be accompanied by visible phenotypic changes in the tumour microenvironment.

Thus, we demonstrate how HistoMIL can be used to assess the detection of thousands of disease-relevant molecules in a speedy and efficient manner, with most informative results obtained for genes that are either expressed throughout the entire slide or not expressed at all within the tissue.

## DISCUSSION

Implementing a multiple instance learning pipeline for WSI datasets requires extensive engineering work and deep understanding of model hyperparameters. The complex nature of WSIs means that extra effort is required in reading, pre-processing and normalisation. In contrast to image processing models employed for conventional classification tasks, whole slide images (WSI) necessitate specialised data retrieval and preprocessing to facilitate deep learning model processing. Concurrently, since H&E slides are relatively sensitive to colour variations, they require an additional normalisation step and should carefully select data augmentation steps (i.e. must not undergo extreme cropping operations). In this paper, we introduce HistoMIL, a Python library for pre- and post-processing WSIs within a MIL-based pipeline. It is designed to work closely with PyTorch, a widely used deep learning toolkit, to allow users to train a batch of MIL-models for various targets easily. The use of PyTorch Lightning simplifies the training process. Also, our package HistoMIL offers a simple way to train a feature extractor with an SSL protocol, which may increase the performance of different methods in the target data domain. We provide a detailed tutorial on how to use this package for both technical and non-technical users in our *Notebooks* folder at https://github.com/secrierlab/HistoMIL.

HistoMIL aims to facilitate the training and application of MIL algorithms for predicting molecular labels based on digital pathology slides by providing an easy-to-use API. In doing so, we hope to provide users a comprehensive set of tools that will enable them to focus on developing new algorithms and addressing biological questions. To achieve this goal, we have implemented a wide range of modules and functions to streamline the process of training and using SSL and MIL. Using HistoMIL’s SSL component, we can effortlessly generate a feature extractor for the diagnostic slide dataset, which can be then used to predict different molecular labels. In our experiments we exemplify the prediction of the expression status of a variety of genes in H&E slides using the built-in MIL algorithms. These pipelines have been implemented in the form of interactive notebooks and can be opened and evaluated on cloud platforms such as Google Colab and Kaggle. This highlights how HistoMIL can be used to greatly simplify the complexity of engineering implementations. We hope that the examples provided will help other users integrate MIL methods as an effective tool in their analysis pipelines.

By designing HistoMIL to maintain consistency and ease of use when introducing customised models, we observed that the data processing steps of different algorithms can be shared due to the repeated use of HistoMIL modules. Additionally, different algorithms can also share the same intermediate data results. Furthermore, batch processing and patch aggregation in HistoMIL come with preset values, which greatly reduces the level of difficulty for users. Here, we emphasise that HistoMIL is not limited to the implemented MIL algorithms. Due to its modular and scalable design, users can conveniently implement and modify new algorithms. This helps in training new customizable algorithms based on existing work. Therefore, any algorithm implementation involving MIL and WSIs can benefit from the functionality we provide. In addition, a large number of tasks involving WSIs can also be conveniently implemented through modifications to algorithms or loss functions.

We demonstrated good performance of HistoMIL on more than 2,000 gene expression prediction tasks using a variety of MIL algorithms, including AUCs above 80% for 130 genes. We showed that amongst classical cancer hallmark pathways the most identifiable within H&E-stained slides are the ones related to cell proliferation, in line with other findings in the field (43), whereas kinase signalling, apoptosis and oxidative stress processes are more difficult to capture from the tissue morphology. This suggests that cancer diagnosis and progression tasks could be more easily automated on histopathology slides than tasks related to targeted treatment decisions. Furthermore, the spatial visualisation capabilities of HistoMIL pave the way towards further analyses of spatial patterns of gene activity, which could be used to understand how cancers develop within the tissue and interact with their environment.

HistoMIL is an open-source project that will continue to add additional pre-trained models and functionality. In the future, we plan to expand the currently available models by training on new datasets and adding other MIL algorithms. A reasonable extension is to apply it to cancer grading, survival prediction and other molecular or clinical labels in different tumour types. However, the package is not limited to analysing cancer datasets and could be easily employed for any biomedical question aiming to identify meaningful patterns in H&E stained slides.

## LIMITATIONS OF THE STUDY

Our experiments revealed some limitations of the existing MIL algorithms. Firstly, as expected, their performance is highly dependent on the image quality of the slides. As the expression levels of some genes are correlated with the colour patterns of the images, the models are sensitive to the colour changes in the original slides. MIL algorithms also incorporate attention mechanisms and neighbourhood information, which makes them more sensitive to variations within a single slide, such as local colour changes. Moreover, patch-level predictions can introduce biases. Some genes might not be expressed in all regions of the slide, but because a single expression label is available per slide, the models would tend to assume high expression in all areas when this label is high.

The number of samples in the tested dataset also limits our analyses. In the context of Transfer Learning, the number of training samples is determined by the number of Whole Slide Images (WSIs) in the dataset multiplied by the number of patches within each WSI. Essentially, each patch extracted from a WSI is treated as an independent sample, leading to a large number of training samples. However, when employing the Multiple Instance Learning method, the number of training samples for the slide-level classifier is equivalent to the number of WSIs in the dataset. This is because in MIL each WSI is treated as a single sample, rather than considering each patch within the WSI as an individual sample. This fundamental difference could potentially impact the generalisation capability of the MIL method. This also makes the model aggregation part prone to being misled by some samples and thus trapped in local optima. Our future work will try to address those problems by expanding the pool of diagnostic slides using other datasets.

## ACKNOWLEDGEMENTS

MS and SP were supported by a UKRI Future Leaders Fellowship (MR/T042184/1). Work in MS’s lab was supported by a BBSRC equipment grant (BB/R01356X/1) and a Wellcome Institutional Strategic Support Fund (204841/Z/16/Z).

We would like to thank Eloise Withnell for testing and providing feedback on the HistoMIL package.

## AUTHOR CONTRIBUTIONS

SP developed the HistoMIL package and performed all the analyses. MS acquired the funding, supervised the analyses and helped interpret the results. Both authors wrote and approved the manuscript.

## DECLARATION OF INTERESTS

The authors have no conflicts of interest to declare.

## ETHICS APPROVAL AND CONSENT TO PARTICIPATE

All data employed in this study comply with ethical regulations, with approval and informed consent for collection and sharing already obtained by The Cancer Genome Atlas (TCGA).

## DATA AND CODE AVAILABILITY

The results published here are based upon publicly available data generated by the TCGA Research Network: https://www.cancer.gov/tcga. The HistoMIL package is available at the following repository: https://github.com/secrierlab/HistoMIL

## Notes

https://github.com/secrierlab/HistoMIL

### Competing Interest Statement

The authors have declared no competing interest.

